# Comparative Genomic Analysis of Metal-Tolerant Bacteria Reveals Significant Differences in Metal Adaptation Strategies

**DOI:** 10.1101/2024.07.03.601927

**Authors:** Dai Di Chen, Liu Lian Zhang, Jiu Hua Zhang, Wen Ting Ban, Qingxin Li, Jin Chuan Wu

## Abstract

Metal-tolerant bacteria have been commercially used in wastewater treatment, bio-fertilizer and soil remediation etc. However, their action mechanisms have not yet been well understood. We screened metal-tolerant bacteria isolated from the rhizosphere soil samples with metal-enriched media containing Cu, Fe or Mn, sequenced and compared the genomes, and analyzed their metal adaptation strategies at genomic levels to better understand their mechanisms of actions. Totally 32 metal-tolerant isolates were identified and classified into 12 genera based on phylogenetic analysis. The determination of MTC and effect of metal ions on the isolates indicated that *Serratia marcescens* X1, *Mammaliicoccus sciuri* X26 and *Rummeliibacillus pycnus* X33 showed the significant differences in metal tolerance to Cu, Fe and Mn with other isolates. They have quite different genomic features to adapt various metal ions. *S. marcescens* X1 possesses abundant genes required for Cu, Fe and Mn homeostasis. *M. sciuri* X26 has a number of genes involved in Mn and Zn homeostasis but with no genes responsible for Cu and Ca transport. *R. pycnus* X33 is rich in Fe, Zn and Mg transport systems but poor in Cu and Mn transport systems. It is thus inferred that the combined use of them would compensate their differences and enhance their ability in accumulating a wider range of heavy metals for promoting their applications in wastewater treatment, soil remediation and organic fertilizer etc.

**IMPORTANCE:** Metal-tolerant bacteria have wide applications in environment, agriculture and ecology, but their action strategies have not yet been well understood. We isolated 32 metal-tolerant bacteria from the rhizosphere soil samples. Among them, *S. marcescens* X1, *M. sciuri* X26 and *R. pycnus* X33 showed the significant differences in metal tolerance to Cu, Fe and Mn with other isolates. Comparative genomic analysis revealed that they have abundant and different genomic features to adapt various metal ions. It is thus inferred that the combined use of them would compensate their differences and enhance their ability in accumulating heavy metal ions widening their applications in industry, agriculture and ecology.

## INTRODCTION

Metal-tolerant bacteria that exhibit high resistance to metal ions, have been found in varied environments and habitats such as water, soil, plants and humans/animals (1–4). Metal-tolerant bacteria possess various metal transport systems to facilitate them as intermediate storage stations for metal ions to shuttle through various environments and habitats. Therefore, metal-tolerant bacteria have been applied in many fields such as treatment of waste liquors, transport of metal ions between plants and soil, regulation of soil pH and remediation of contaminated soil (5–8). With the increasing roles played by metal-tolerant bacteria in human life, recently multi-omics technologies (e.g., genomics, transcriptomics, proteomics and metabolomics) have been applied for investigating the metal resistance mechanisms of microorganisms (9–12).

Metal ions are required as essential nutrients for survival of organisms because they play important roles in cellular metabolisms. It is noteworthy that excessive metal ions can have significantly harmful effects on living organisms (13, 14). To survive in environments of varied metal concentrations, microorganisms need diverse strategies to maintain intracellular metal homeostasis. Studies on mechanisms of metal homeostasis in microorganisms suggested that metal transport systems and metal-binding proteins are key players in response to metal stress (15, 16). In bacteria, metal transport systems are conserved but diverse, such as <u>A</u>TP-<u>b</u>inding <u>c</u>assette (ABC) transporters (e.g., *<u>A</u>ctinobacillus* <u>f</u>erric <u>u</u>ptake system AfuABC required for iron (Fe) influx (17), *<u>S</u>almonella* iron transporter SitABCD involved in the acquisition of Fe and manganese (Mn) (18), <u>Mn</u> transporter MntABC responsible for Mn uptake (19) and zinc (<u>Zn</u>) <u>up</u>take system ZnuABC necessary for Zn import (15)), two-component system (e.g., CusSR and PcoSR regulating copper (Cu) influx (20), PhoPQ mediating magnesium (Mg) import (21) and potassium (<u>K</u>)-<u>d</u>ependent <u>p</u>rotein KdpDE controlling K uptake (22)), <u>n</u>atural <u>r</u>esistance-<u>a</u>ssociated <u>m</u>acrophage <u>p</u>roteins (NRAMP) (e.g., <u>Mn</u> transporter MntH involved in Mn import (16)), P-type ATPases (e.g., CopA contributing to the Cu efflux (20)), symporter (e.g., glutamate:Na^+^ symporter GltS, solute:Na^+^ symporter SSS and proline:Na^+^ symporter PutP performing sodium (Na) influx (23)), and antiporter (e.g., Na^+^:H^+^ antiporter NhaABC and multicomponent Na^+^:H^+^ antiporter MnhA-G carrying out Na efflux (24), and <u>C</u>a^2+^:<u>H</u>^+^ <u>a</u>ntiporter ChaA undertaking calcium (Ca) export (25)).

In order to obtain metal-tolerant bacteria, we screened the isolates from rhizosphere soil samples with metal-enriched medium containing Cu, Fe or Mn, and performed phylogenetic analysis based on the16S rRNA sequences of the isolates. Moreover, we sequenced and compared their genomes, analyzed their metal adaptation strategies at genomic level to provide insight into their differences in action mechanisms for guiding the development of their new applications.

## RESULTS AND DISCUSSIONS

### Isolation and molecular identification of metal-tolerant bacteria

Nearly 100 isolates from rhizosphere soil samples were screened using the metal-enriched medium containing 200 mg/L each of CuSO_4_, FeSO_4_ or MnSO_4_·4H_2_O. Totally 32 isolates were identified with 16S rRNA gene sequences revealing their phylogenetic affiliation with 12 genera, including *Pseudomonas*, 8; *Cupriavidus*, 5; *Enterobacter*, 4; *Klebsiella*, 4; *Bacillus*, 3; *Chryseobacterium*, 2; *Atlantibacter*, 1; *Lelliottia*, 1; *Exiguobacterium*, 1; *Rummeliibacillus*, 1; *Serratia*, 1; *Mammaliicoccus* 1. The Fe-, Mn- and/or Pb-tolerant bacteria, which were isolated from coal mine samples, belong to the genera of *Bacillus*, *Serratia*, *Rhodococcus* and *Paenibacillus* (3). Cidre et al. (2) isolated the Cu- and Zn-resistant bacteria from fresh produces and identified them to 6 genera: *Enterobacter*, *Leclercia*, *Pseudomonas*, *Serratia*, *Bacillus* and *Paenibacillus*. These results suggest that the species of *Enterobacter*, *Pseudomonas*, *Serratia* and *Bacillus* might act as the key players in transformation of metal (Fe, Cu or/and Mn) tolerance in soil and plants.

Among the 32 isolates, 16 of them exhibited distinct colony morphologies and the ones that were at least tolerant to two metals, were selected for further investigation (Fig. 1). The phylogenetic relationships of the 16 isolates and the related species were analyzed using neighbor-joining phylogenetic tree with their 16S rRNA gene sequences (Fig. 2; Table 1). The 16S rRNA gene identity > 98% is a traditional microbial species delineation to estimate two strains being the same species (37). The isolates X1, X3 and X5 showed 16S rRNA gene sequence similarities of 99.72%, 99.65% and 99.93% with *Serratia marcescens* C3, *Lelliottia jeotgali* N1 and *Atlantibacter hermannii* R11, respectively, so the isolates X1, X3 and X5 were identified as *S. marcescens*, *L. jeotgali* and *A. hermannii*. The isolates X4 and X17 were homologous to *hryseobacterium* spp. (> 98.43% similarities), so they were classified as *C. gleum* and *C. defluvii*, respectively. The isolates X6 and X12 clustered together with *Enterobacter* spp. by > 99.72% of 16S rRNA gene identities, so they were identified as *E. cloacae* and *E. ludwigii*, respectively. The isolates X8, X11 and G2 were most matching with *Bacillus* spp. with 16S rRNA gene sequence similarities of > 99.79%, so they were designated as *B. pseudomycoides* X8, *P. megaterium* X11 and *B. subtilis* G2, respectively. The 16S rRNA gene sequences of isolates X10 and N4 were 99.79% and 99.38% similar to those of *Exiguobacterium indicum* MnW2201005 and *Cupriavidus gilardii* CR3, so they were classified as *E. indicum* and *C. gilardii*, respectively. The16S rRNA gene sequences of isolates X19 and X30 correlated well with those of *Pseudomonas* spp. (> 99.72%), so they were identified as *P. plecoglossicida* and *P. nitroreducens*, respectively. The isolates X26 and X33 were closest to *Mammaliicoccus sciuri* CB212 and *Rummeliibacillus pycnus* 111_12 with great bootstrap supports (100% and 99%) and 16S rRNA gene sequence similarities (99.72% and 99.08%, respectively), Therefore, they were designated as *M. sciuri* X26 and *R. pycnus* X33, respectively.

**Fig. 1.**
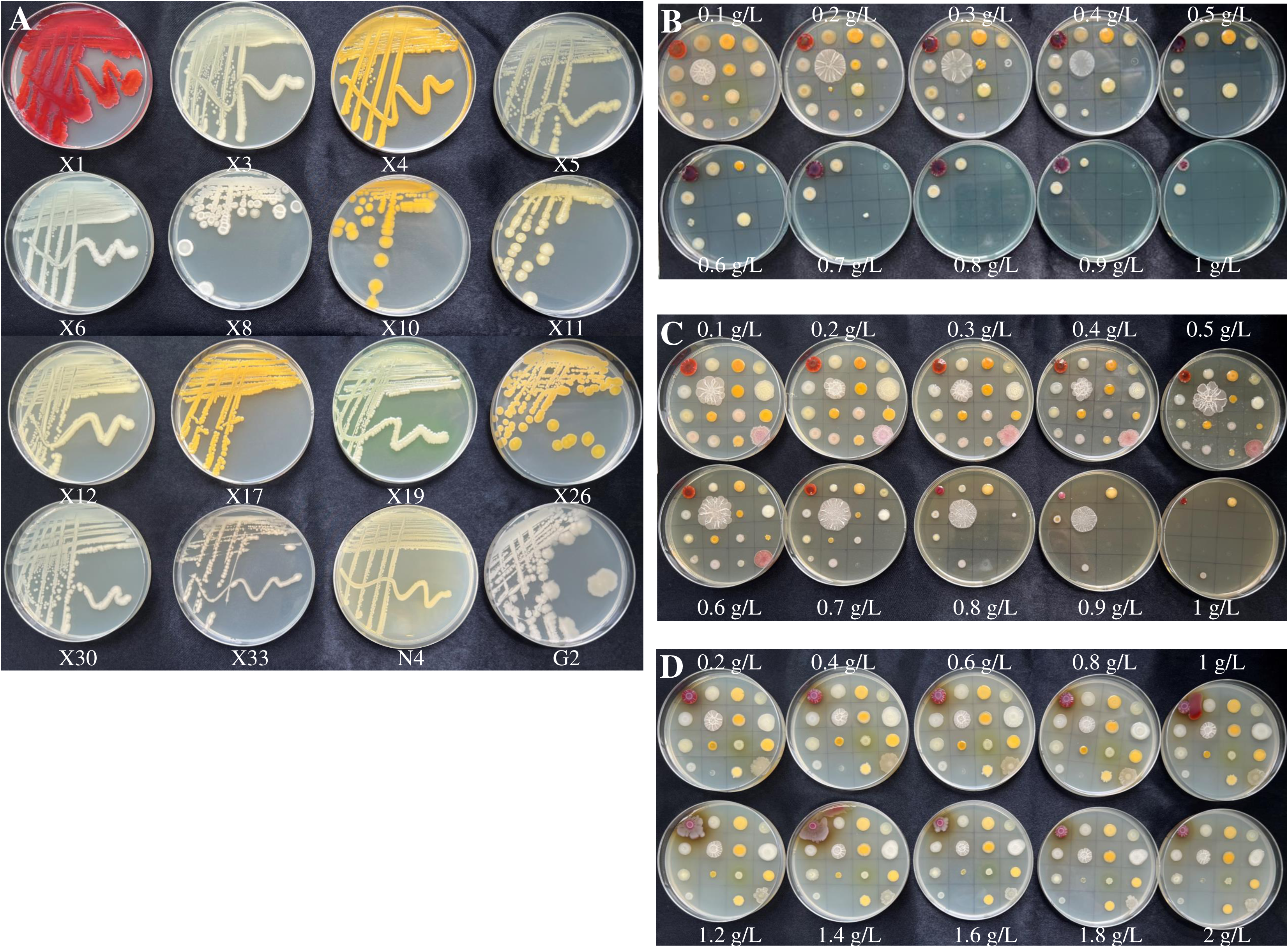
MTC of 16 metal-tolerant bacterial isolates. 16 isolates were grown on LB medium (A). 16 isolates were grown on LB medium supplemented with various concentrations of CuSO_4_ (B), FeSO_4_ (C) and MnSO_4_·4H_2_O (D). Photographs were taken 3 days after inoculation at 30°С.

**Fig. 2.**
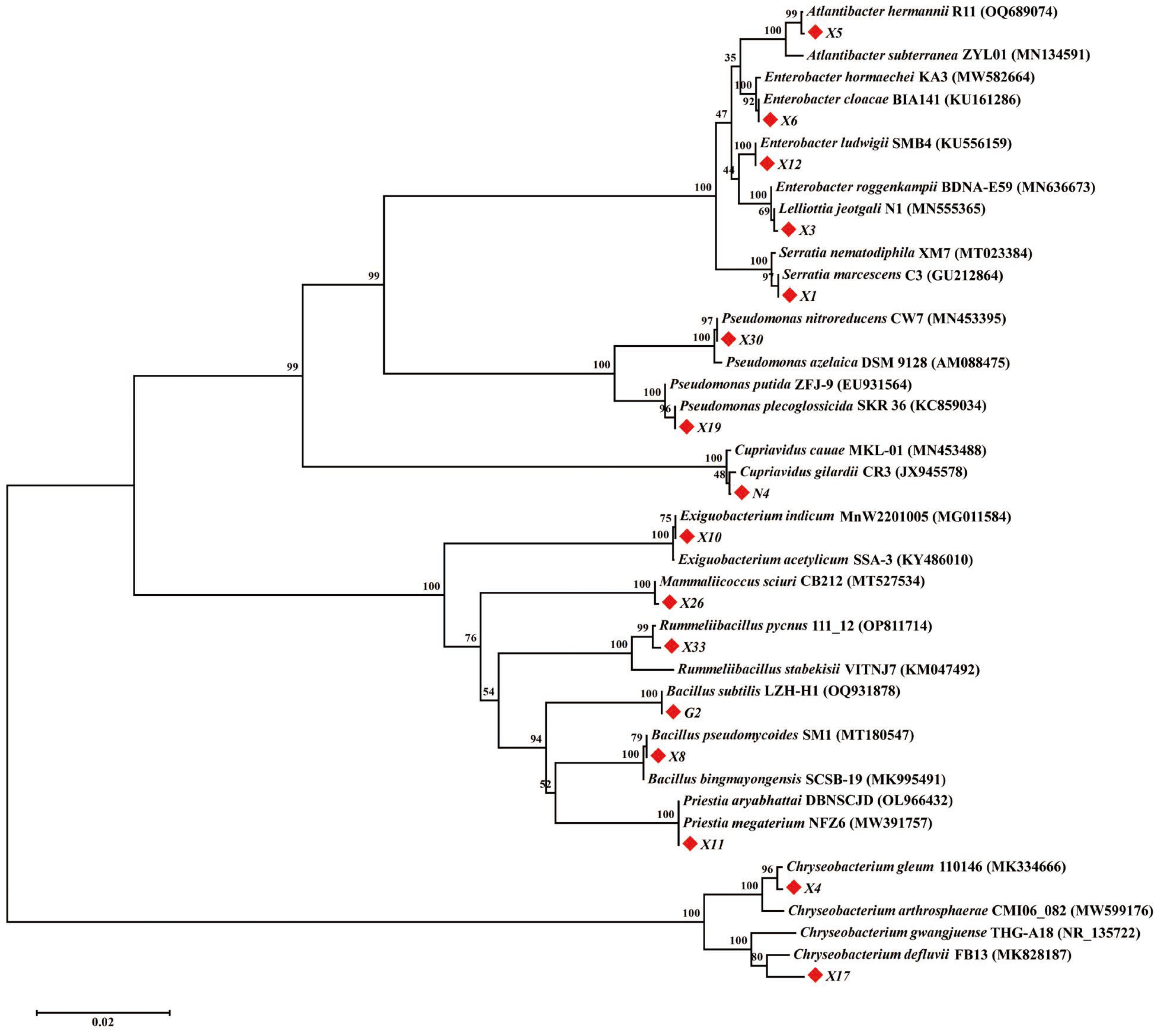
Phylogenetic relationships of 16 metal-tolerant bacterial isolates and the related species. Neighbour-joining phylogenetic tree based on 16S rRNA gene sequences showing the phylogenetic positions of 16 metal-tolerant bacterial isolates (indicated with solid red diamond). The bootstrap values are shown at branch nodes. Bar, 0.02 nucleotide position.

**Table 1.**
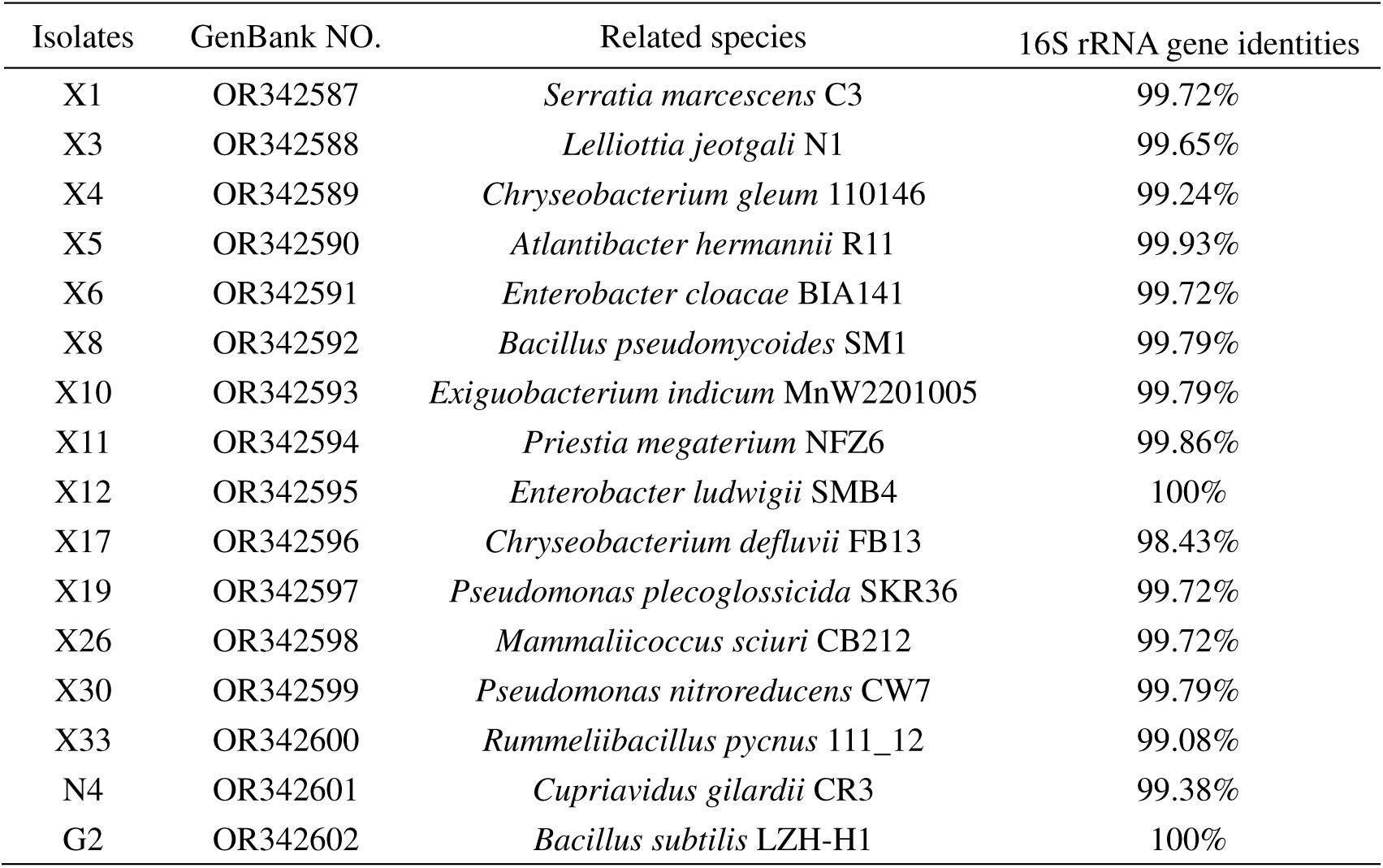
16S rRNA gene identities of metal-tolerant bacterial isolates with Related species.

### Maximum tolerance concentration (MTC) of metal-tolerant bacteria

The MTC of the 16 isolates for Cu, Fe and Mn was determined using the spot plate method and the results were shown in Fig 1B-D. For Cu, the highest MTC (1000 mg/L) was exhibited by *S. marcescens* X1 and *E. cloacae* X6, followed by *L. jeotgali* X3 and *A. hermannii* X5 (900 mg/L). Besides, all isolates could tolerate a minimum concentration of 200 mg/L of CuSO_4_, except for *M. sciuri* X26. In the case of Fe, *S. marcescens* X1, *C. gleum* X4 and *R. pycnus* X33 displayed the highest MTC of 1000 mg/L, accompanied by *E. cloacae* X6 and *B. pseudomycoides* X8 (MTC of 900 mg/L). Moreover, all isolates could tolerate a minimum concentration of 600 mg/L of FeSO_4_. All isolates could grow at a concentration of 2000 mg/L of MnSO_4_·4H_2_O, except for *R. pycnus* X33, which showed a MTC of 400 mg/L against Mn. Overall, *S. marcescens* X1 exhibited the highest MTC for Cu, Fe and Mn, *M. sciuri* X26 showed the highest MTC for Mn and the lowest MTC for Cu, and *R. pycnus* X33 displayed the highest MTC for Fe and the lowest MTC for Mn. As the above three isolates showed significant differences with others, they were selected for further investigation.

*Enterobacterales* have been reported existing in multiple niches including soil, water, plants, animals and humans. Many species of this order have exhibited high resistance to multiple metals and antibiotics (2–4). It is worth noting that the isolates belonging to the *Enterobacterales* order (*S. marcescens* X1, *L. jeotgali* X3, *A. hermannii* X5 and *E. cloacae* X6) exhibited superior tolerance to multiple metals. The isolate *S. marcescens* X1 showed higher tolerance to metals than other reported *S. marcescens* stains such as *S. marcescens* KH-CC (MTC of 500 ppm against Fe and 830 ppm against Mn) (3), *S. marcescens* CL11 and *S. marcescens* CL35 tolerated 1200 mg/L of Mn (38). Shylla et al. (3) reported that *B. pseudomycoides* KH-12A showed 400 ppm tolerance to Fe and 730 ppm to Mn, obviously lower than those of *B. pseudomycoides* X8. The MTC values of 90-1000 ppm against Fe (3, 39) and MTC of 120-800 ppm against Mn (3) have been reported in other *Bacillus* species, while the isolates *P. megaterium* X11 tolerated 800 mg/L of Fe and 2000 mg/L of Mn, and *B. subtilis* G2 resisted 600 mg/L of Fe and 2000 mg/L of Mn.

### Effect of metals on bacterial growth

The growth kinetics of the isolates X1, X26 and X33 cultured on LB broth with middle concentrations of metals were monitored to investigate the effects of metals (Cu, Fe and Mn) on bacterial growth. A nonsignificant difference was observed in the growth curves of isolate X1 at non-metals, 500 mg/L of CuSO_4_, 500 mg/L of FeSO_4_ and 1000 mg/L of MnSO_4_·4H_2_O, reaching an OD_600_ of approximately 1.9 after 30 h (Fig. 3A), indicating that the middle concentrations of Cu, Fe and Mn for isolate X1 did not obviously affect the growth of this isolate. As the isolate X26 could not resist Cu which was evidenced from the result of MTC test, the effect of Cu was not carried out here. Besides, the isolate X26 reached OD_600_ values of 3.41, 3.34 and 2.81 at non-metals, 300 mg/L of FeSO_4_ and 1000 mg/L of MnSO_4_·4H_2_O after 30 h, respectively (Fig. 3B), suggesting that the middle concentration of Fe for isolate X26 slightly inhibited its growth, while the middle concentration of Mn for isolate X26 significantly inhibited its growth. The isolate X33 grew in LB broth containing 500 mg/L of FeSO_4_ after 30 h giving an OD_600_ of 1.48, which is close to the case of the absence of metals (OD_600_=1.55) (Fig. 3C). In contrast, the isolate got lower growth rates with OD_600_ values of 0.76 and 1.04 at 300 mg/L of CuSO_4_ and 1000 mg/L of MnSO_4_·4H_2_O after 30 h, respectively (Fig. 3C). These results suggest that Cu has the strongest inhibitory effect on the growth of isolate X33, followed by Mn. Overall, the isolate X1 could resist various metals at high concentrations, and its metal tolerance ability is the strongest, followed by isolate X33 then isolate X26. In addition, Cu is most toxic to the isolates, followed by Mn then Fe.

**Fig. 3.**
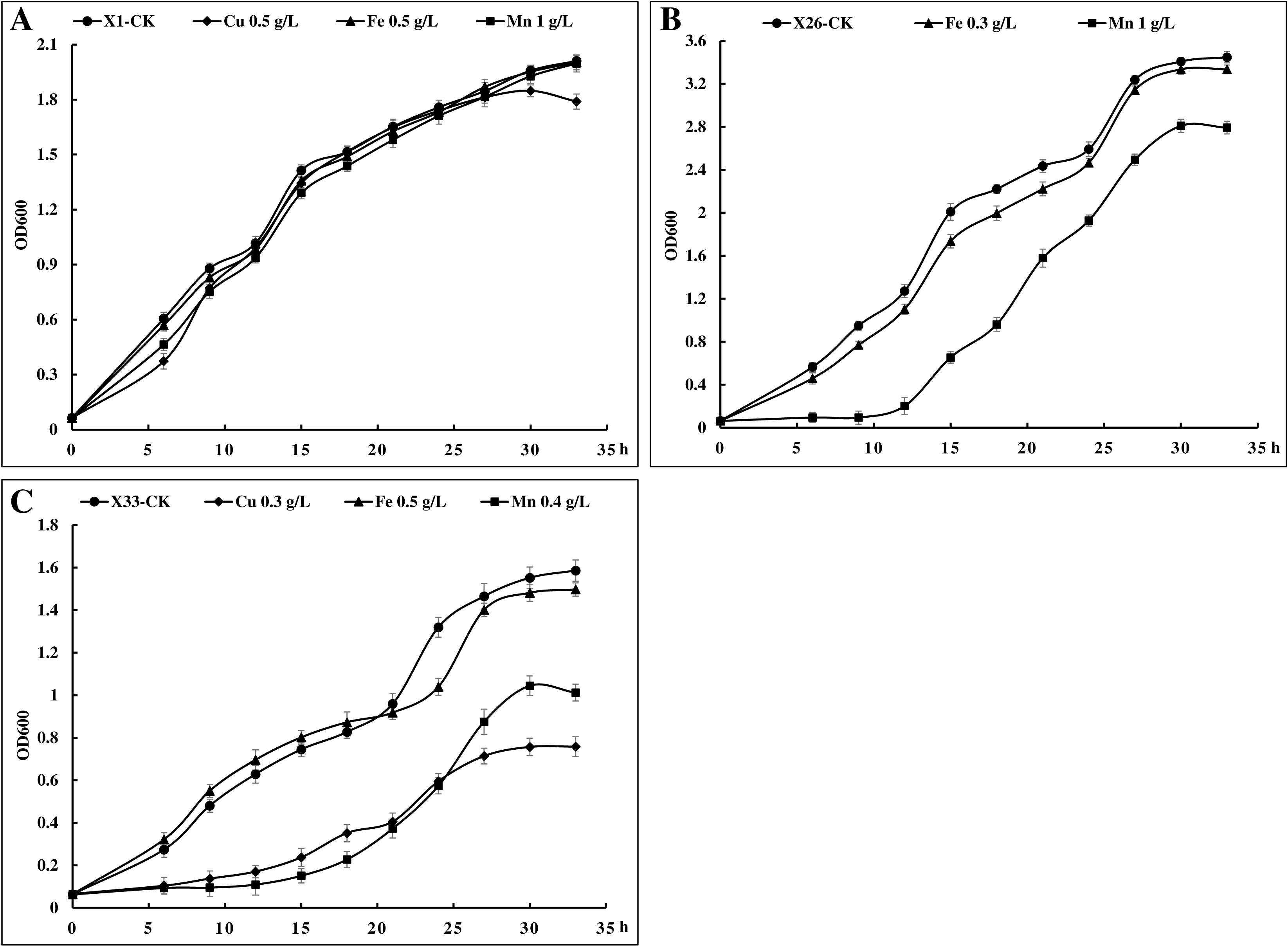
Effect of metals on bacterial growth. *S. marcescens* X1 (A), *M. sciuri* X26 (B) and *R. pycnus* X33 (C) in presence of Cu, Fe and Mn.

### Genomic characteristics of the three metal-tolerant bacterial isolates

Genomic features of the three metal-tolerant bacterial isolates are shown in Fig. 4 and Table 2. The complete genome of *S. marcescens* X1 is assembled in a single circular chromosome of 4,962,287 bp (G+C content of 59.82%), which is smaller than that of other *S. marcescens* strains such as N2 (5.14 Mbp), RSC-14 (5.13 Mbp) and WW4 (5.24 Mbp) (10, 40). A total of 4,542 genes were predicted from the genome of *S. marcescens* X1, including 92 tRNA genes and 22 rRNA genes. The draft genome of *M. sciuri* X26 (3,117,130 bp with a G+C content of 32.56%) assembly contains a complete circular chromosome (2,982,601 bp) and two plasmids (Plasmid I: 67,732 bp and Plasmid II: 66,797 bp). There are 3,085 genes in the genome, which include 58 tRNA genes and 19 rRNA genes. Compared with *M. sciuri* X26, *M. sciuri* IMDO-S72 has a smaller size of genome (2,898,343 bp) containing a chromosome and four plasmids and fewer genes (2,923) (41). The final genome of *R. pycnus* X33 is a complete circular chromosome with a size of 4,346,358 bp and a GC content of 34.97%, which is bigger than that of *R. pycnus* DSM 15030 (3,851,193 bp) (NJAS01000000). Totally, 4,105 predicted genes were found in the genome of *R. pycnus* X33, of which 122 are tRNA genes and 39 are rRNA genes. In addition, 4,532, 3,046 and 3,971 coding sequences (CDSs) were functionally annotated with COG, GO, Swiss-Prot, NR and KEGG database from the genome of *S. marcescens* X1, *M. sciuri* X26 and *R. pycnus* X33, respectively (Supplementary Table S1). Functional categories of genes based on COG and KEGG database are shown in Supplementary (Fig. S1).

**Fig. 4.**
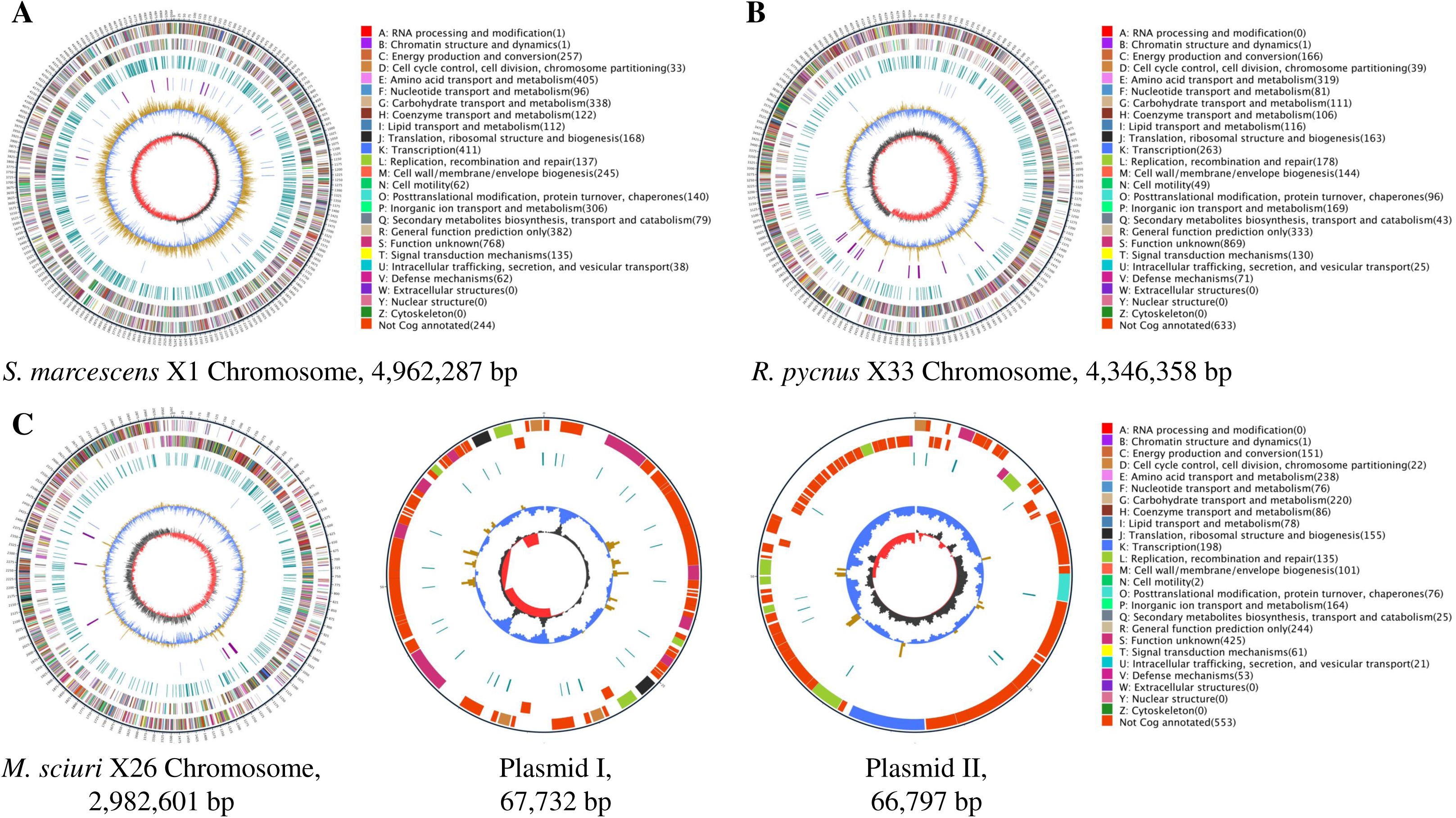
Circular maps of *S. marcescens* X1 (A), *R. pycnus* X33 (B) and *M. sciuri* X26 (C) genomes. From inner to outer circles: (1) GC skew. Dark gray represents areas with G content greater than C, while red represents areas with C content greater than G. (2) GC content. The value is plotted as the deviation from the average GC content of the entire sequence. (3) tRNA (blue) and rRNA (purple). (4) Repetitive sequences. (5, 6) CDS coloured according to COG functional categories, 5 is negative strand, 6 is positive strand. (7) Marker of genome size, with each scale measuring 5 kb.

**Table 2.**
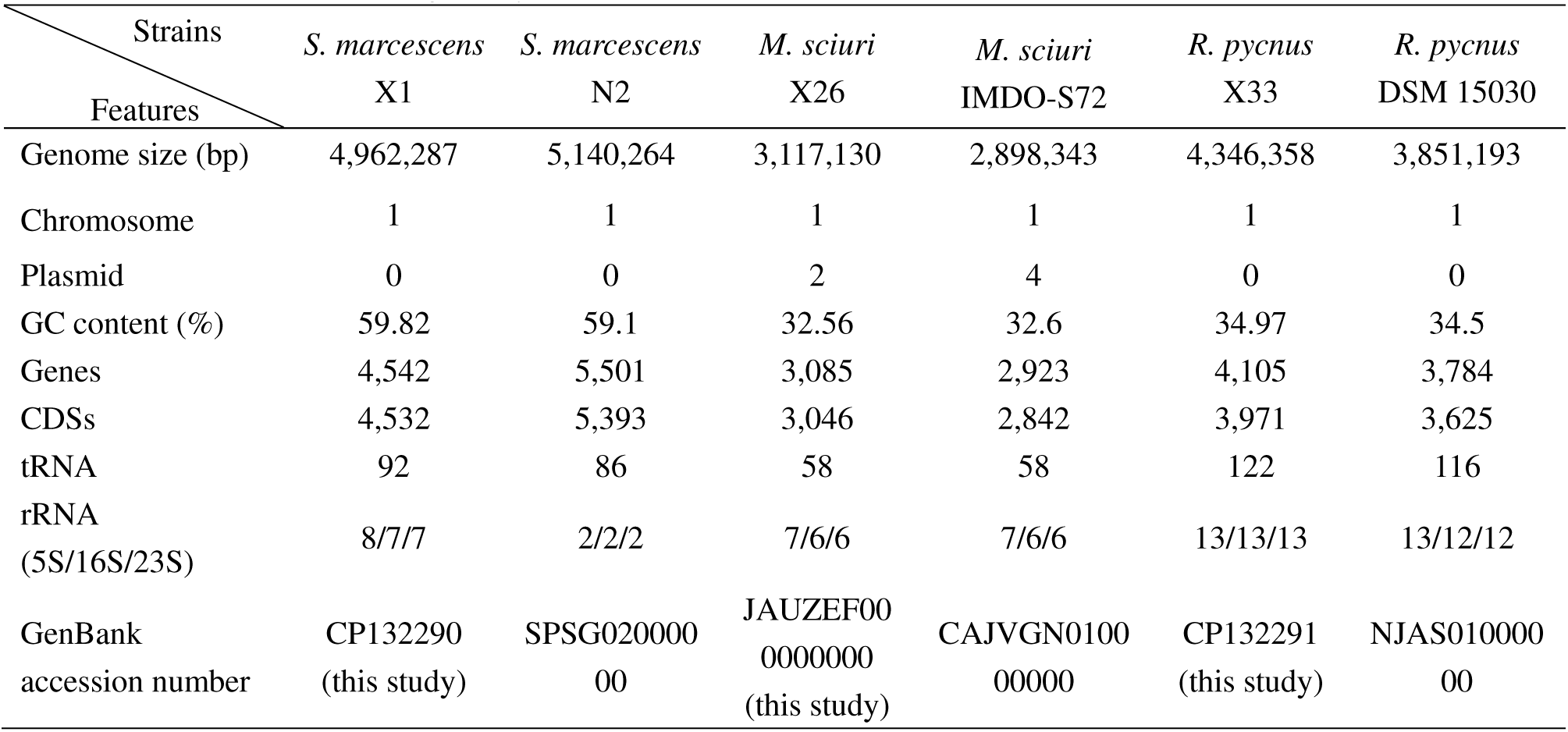
Comparative genome statistics of three metal-tolerant bacterial isolates and other strains of homogeneity.

### Metal adaptation strategies of the three metal-tolerant bacterial isolates

Metal ions are essential micronutrients for biological systems, forming active centers of various metalloenzymes. However, excess amount of metals is cytotoxic. Therefore, elucidation of the mechanisms of metal responses in bacteria would help develop sophisticated cultivation strategies for new applications. To investigate the metal adaptation strategies of metal-tolerant bacteria at a genomic level, the genes responsible for metal tolerance pathways in the genomes of the three metal-tolerant bacterial isolates were identified and compared (Fig.5 and Supplementary Table S4). It is seen that the putative metal stress-response genes are different in both types and amounts, which likely results in their different metal adaptation strategies.

**Fig. 5.**
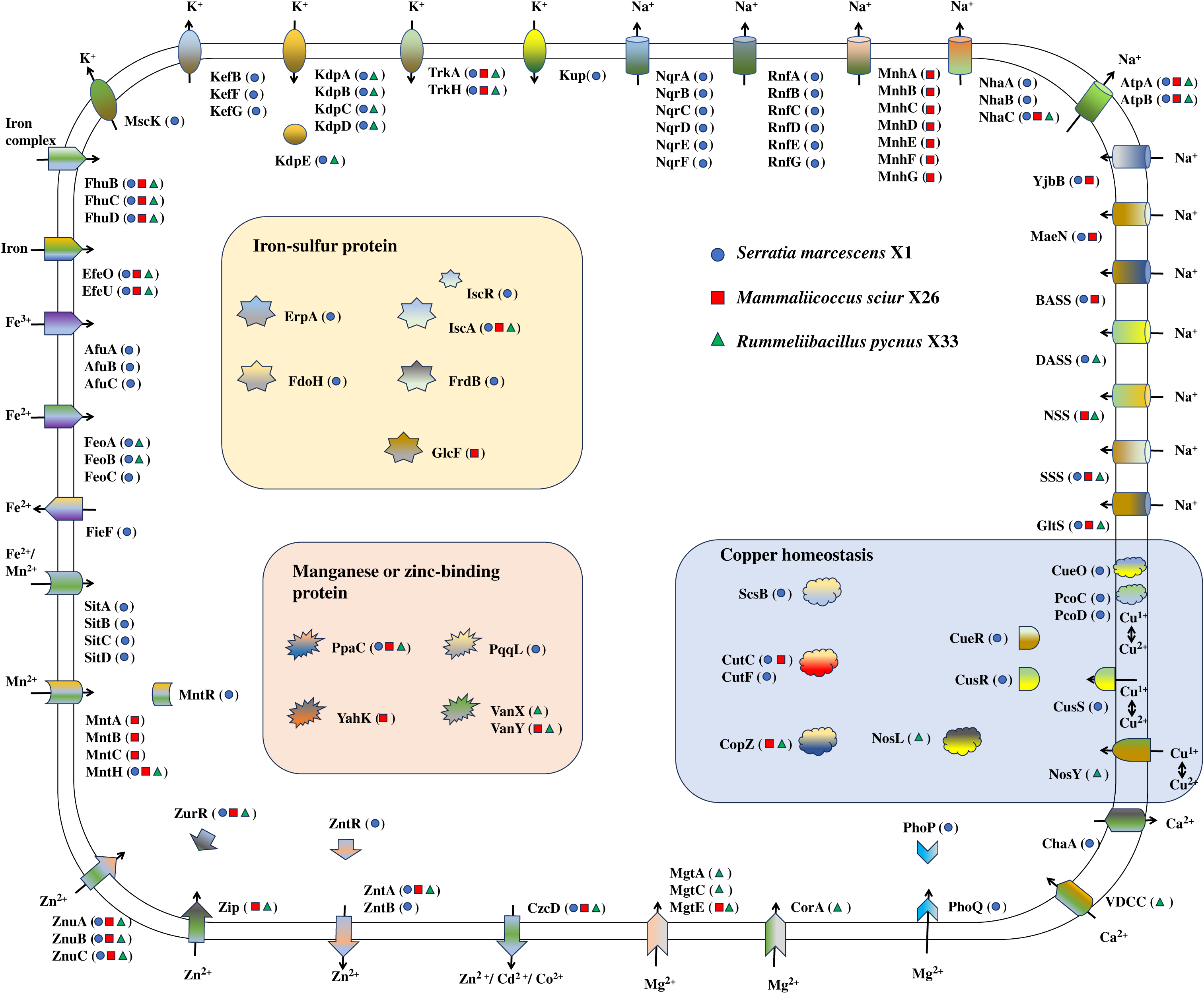
Schematic of the proteins identified as playing a putative role in the adaptation of three metal-tolerant bacterial isolates to metal environments. Strains: *S. marcescens* X1 (blue solid circle), *M. sciuri* X26 (red solid square) and *R. pycnus* X33 (green solid triangle). The putative function of genes is shown in Supplementary Table S2.

For copper homeostasis, in *S. marcescens* X1, the homologs of a multi-copper oxidase, CueO, and two periplasmic chelators, PcoC and PcoD, probably perform copper detoxification in the periplasm by converting periplasmic Cu(I) to Cu(II) (42, 43). The Cu(I) in periplasm fluxes across the inner membrane under the control of the homologs of three sensory systems, i.e., the MerR-family transcriptional regulator (CueR) and the CusS/CusR two-component system (20). The Cu(I) in cytoplasm may be scavenged by the homologs of a high-affinity cuprochaperone (CopZ) and transported by two copper homeostasis proteins (CutC and CutF) (20, 44). *S. marcescens* X1 shows high copper resistance most likely due to the rich of genes involved in copper homeostasis in its genomes. *M. sciuri* X26 possesses CopZ and CutC but lacks the copper sensory systems, likely resulting in conditional lethality in the presence of exogenous copper. The copper tolerance of *R. pycnus* X33 (MTC of 400 mg/L) was lower than *S. marcescens* X1 (MTC of 1000 mg/L), likely owing to the poorer copper response systems, i.e., two copper chaperone (CopZ and NosL) and one Cu-processing system permease protein (NosY).

In *S. marcescens* X1 genome, a number of genes encoding the homologs of proteins responsible for Fe uptake pathways were discovered, including the <u>f</u>erric <u>h</u>ydroxamate <u>u</u>ptake (Fhu) system (FhuB, -C and -D) necessary for the transport of ferrichrome, and other Fe^3+^-compounds from the periplasm (45), <u>e</u>lemental <u>Fe</u>(II/III) (Efe) complex (EfeO and EfeU) required for the acquisition and utilization of iron (46), the AfuABC system responsible for the uptake of Fe(III) (17), the <u>fe</u>rr<u>o</u>us iron transport (Feo) system (FeoA, -B and -C) involved in the transport of Fe(II) (47), and the *<u>S</u>almonella* iron transporter (Sit) (SitA, -B, -C and -D) which plays an important role in Fe acquisition under iron-deficient conditions (48). Many iron-sulfur proteins, i.e., iron-sulfur cluster assembly protein (IscA)/iron-sulfur cluster assembly regulator (IscR), A-type iron-sulfur carrier protein (ErpA), fumarate reductase iron-sulfur subunit (FrdB), and formate dehydrogenase iron-sulfur subunit (FdoH), have been reported to play important roles in Fe homeostasis and many cellular processes (49–52). Therefore, the determination of MTC and effect of iron on *S. marcescens* X1 show that this isolate exhibited high tolerance to iron (MTC of 1000 mg/L) and its growth was not obviously affected by middle concentration of iron (500 mg/L of FeSO_4_). *R. pycnus* X33 also exhibited an MTC of 1000 mg/L against Fe, but its growth obviously decreased under 500 mg/L FeSO_4_, which might be ascribed to the absence of some Fe uptake systems (i.e., AfuABC system and SitABCD transporters) and iron-sulfur proteins (ErpA, FrdB and FdoH) in *R. pycnus* X33 compared with *S. marcescens* X1. *M. sciuri* X26 appears to absorb iron only via two Fe uptake systems, FhuBCD system and EfeUO complex, resulting in the lower copper tolerance than other two isolates.

*S. marcescens* X1 exhibited high tolerance to Mn (MTC of 2000 mg/L), possibly due to the two types of Mn import systems present in this isolate. The first system is a NRAMP family member, MntH, which is a high-affinity manganese acquisition system regulated by MntR (16, 18). The second system is an ABC transporter, SitABCD, which transports Mn also via the mediation of MntR (18). *M. sciuri* X26 showed high tolerance to Mn (MTC of 2000 mg/L), possibly resulting from two manganese transporters, i.e., homologs of MntH and ABC transporter MntABC (19). However, the absence of MntR might reduce the Mn tolerance of this isolate, so its growth decreased under 1000 mg/L MnSO_4_·4H_2_O. The Mn tolerance of *R. pycnus* X33 was lowest among the three isolates, which might be attributed to the only one manganese transporter (MntH homolog) in this isolate.

In the genomes of the three isolates, a number of genes involved in the transportation and homeostasis of other metals (i.e., Zn, Mg, Ca, Na and K) were also found. For the transportation of Zn, the Znu and Zip (<u>Z</u>rt/<u>I</u>rt-like <u>p</u>rotein) systems are believed to be responsible for the influx of Zn into the cells, while the Znt (<u>Zn</u> transporter) and Czc (<u>c</u>obalt, <u>z</u>inc and <u>c</u>admium) systems are thought to be involved in the efflux of Zn out of the cells (15, 16, 53). For the Zn uptake, the homologs of *znuABC* and their regulatory gene *zurR* were found in the genomes of the three isolates, while the homolog of *zip* is only present in the genomes of *M. sciuri* X26 and *R. pycnus* X33. For the Zn efflux, *S. marcescens* X1 possesses the homologs of ZntABR and CzcD, while *M. sciuri* X26 and *R. pycnus* X33 only have ZntB and CzcD homologs. Moreover, some genes, *vanX* (encoding zinc-dependent D-Ala-D-Ala dipeptidase), *vanY* (encoding zinc-containing D-Ala-D-Ala carboxypeptidase) and *yahK* (encoding zinc-type alcohol dehydrogenase-like protein), involved in the Zn homeostasis and many physiological processes (54, 55), were found in the genomes of *M. sciuri* X26 and/or *R. pycnus* X33 but not in the genome of *S. marcescens* X1. These findings suggest that *M. sciuri* X26 and *R. pycnus* X33 may have a superior ability for resisting Zn compared to *S. marcescens* X1. The main Mg transport systems of the three isolates show differences. The Mg uptake of *S. marcescens* X1 is likely mediated by the homologs of PhoP (the response regulator/transcriptional activator)/PhoQ (the sensor/receptor histidine kinase) two-component system (21). In *M. sciuri* X26, the Mg import into cells may be performed by transporter MgtE, a widely distributed Mg^2+^ channel in microorganisms (56). The four Mg transporters, CorA, MgtA, MgtB and MgtE, reported to be highly selective for Mg uptake (57), probably contribute to the Mg transport in *R. pycnus* X33. In the case of Ca transport, one homolog of VDCC (<u>v</u>oltage-<u>d</u>ependent <u>C</u>a^2+^-<u>c</u>hannel) involved in the Ca^2+^ influx (58), was only found in *R. pycnus* X33, and one homolog of ChaA responsible for the Ca^2+^ efflux (25), is only present in *S. marcescens* X1.

It is well known that the intracellular concentrations of K and Na are very important to the physiological processes and salt-stress-response in microorganisms. In microorganisms, the K^+^ influx is mainly performed by four systems, the Trk/Ktr (<u>K</u>-<u>tr</u>ansport), Kdp and Kup (<u>K</u>^+^ <u>up</u>take) systems, while the K^+^ efflux is mainly carried out by the glutathione-gated potassium (<u>K</u>) <u>ef</u>flux (Kef) system and <u>m</u>echano<u>s</u>ensitive <u>c</u>hannels (Msc) (23). Among these K^+^ uptake systems, Trk and Kup are constitutive systems, while Kdp is an inducible system, exhibiting high affinity for K^+^ (59). In *S. marcescens* X1, three K^+^ influx systems, KdpABCDE, TrkAH and Kup were found, and in *R. pycnus* X33, two K^+^ uptake systems, KdpABCDE and TrkAH, were discovered, while in *M. sciuri* X26, only TrkAH was observed. It is noteworthy that *S. marcescens* X1 possesses the homologs of KefBFG and MscK for K^+^ efflux, but no homologs of Kef or Msc were found in *M. sciuri* X26 and *R. pycnus* X33. It is likely that *M. sciuri* X26 and *R. pycnus* X33 as well as *S. marcescens* X1 do not have a strong ability for K^+^ uptake, so they require reducing K^+^ efflux to maintain intracellular K homeostasis. In microorganisms, the uptake of Na is performed by the important secondary active transporters (Na^+^-symporters) (23). In the genomes of the three isolates, some genes homologus to Na^+^-symporters, i.e., glutamate:Na^+^ symporter (*gltS*), solute:Na^+^ symporter (*sss*), neurotransmitter:Na^+^ symporter (*nss*), divalent anion:Na^+^ symporter (*dass*), bile acid:Na^+^ symporter (*bass*), malate:Na^+^ symporter (*maeN*) and phosphate:Na^+^ symporter (*yjbB*) were found. Except for the homolog of NSS, the others six Na^+^-symporters were found in *S. marcescens* X1, while only the homolog of *dass* is not present in the genome of *M. sciuri* X26. Correspondingly, in *S. marcescens* X1 and *M. sciuri* X26, there are many systems performing Na^+^ efflux to maintain intracellular Na homeostasis, including Na^+^-translocating NADH:ubiquinone oxidoreductase complex (NqrA-F), Na^+^-translocating NAD^+^:ferredoxin oxidoreductase complex (RnfA-E and RnfG), F-type H^+^:Na^+^-translocating ATPase complex (AtpAB), NhaABC and/or MnhA-G (24, 60–62). Different with *S. marcescens* X1 and *M. sciuri* X26, *R. pycnus* X33 only has four Na^+^-symporters, i.e., GltS, SSS, NSS and DASS, and the Na^+^ efflux is only carried by AtpAB and NhaC.

In summary, 32 metal-tolerant isolates screened from the rhizosphere soil samples with metal-enriched media containing Cu, Fe or Mn, were identified and classified into 12 genera. The determination of MTC and effect of metals on the metal-tolerant isolates indicated that among them, *S. marcescens* X1, *M. sciuri* X26 and *R. pycnus* X33 showed significant differences in metal tolerance to Cu, Fe and Mn with other isolates. *S. marcescens* X1 could resist high concentrations of Cu, Fe and Mn, *M. sciuri* X26 could resist high content of Mn but not Cu, and *R. pycnus* X33 could tolerate high level of Fe but not Mn. To investigate their metal adaptation strategies at a genomic level, the genomes of the three isolates were sequenced and compared. It was found that the genomic features which have been believed to play an important role in adapting metal environments, are significantly different for the three isolates. *S. marcescens* X1 possesses a number of genes encoding transporters, channels and metal-binding proteins to adapt the environments containing high concentrations of Cu, Fe and Mn. *M. sciuri* X26 has a number of genes involved in Mn and Zn homeostasis but no genes responsible for Cu and Ca transport. *R. pycnus* X33 is rich in Fe, Zn and Mg transport systems but poor in Cu and Mn transport systems. Based on these findings, *S. marcescens* X1 is expected to serve as an eco-friendly biosorbent or biological material for effectively removing Cu, Fe and Mn from wastewater and polluted soil, *M. sciuri* X26 is predicted to be used for treating the environment containing high amounts of Mn and Zn, and *R. pycnus* X33 is desired for accumulating Fe, Zn and Mg from environment. The combined use of them might serve as efficient plant growth regulators for the transport of metals between plants and soil as well as for the regulation of soil pH. Additionally, *S. marcescens* X1 might play a role in quick restoration of salt-contaminated soils considering its abundant K and Na transport systems, which were found in halophilic and halotolerant microbes.

## MATERIALS AND METHODS

### Source and isolation of metal-tolerant bacteria

The soil samples were collected from the rhizosphere of banana and sugarcane grown at Guangzhou, Guangdong province, southern China and stored aseptically in a refrigerator at 4°С until processed for isolation. The soil samples (5 g) were diluted to a concentration of 10^-3^ and 10^-4^ (w/v) with autoclaved distilled water, the dilutes were then spread (100 μL per plate) onto modified LB agar plates (HKM, Guangdong, Chain) supplemented with 200 mg/L each of CuSO_4_, FeSO_4_ or MnSO_4_·4H_2_O (Macklin, Shanghai, Chain), and the pH was adjusted to 7.2 ± 0.3. The isolates were selected based on varying colony morphology after 2 d incubation at 30°С, then purified, cultivated and maintained on the same medium. Pure cultures were preserved as 20% (w/v) glycerol at −80°С.

### Phylogenetic analysis of metal-tolerant bacterial isolates

The identification of metal-tolerant bacterial isolates was performed by sequencing the 16S ribosome RNA (rRNA) genes. The 16S rRNA gene was amplified with forward 27F and reverse 1492R primers, and sequenced on an Illumina platform (Illumina NovaSeq 6000) at Tsingke Biotech (Bejing, Chain). The partial sequences of the 16S rRNA gene sequences of the metal-tolerant bacterial isolates were deposited in GenBank under the accession numbers OR342587-OR342602. For phylogenetic analysis, 16S rRNA sequences of type strains with validly published prokaryotic strains affiliated with the isolates were retrieved from the GenBank database. The 16S rRNA gene sequences were aligned to produce phylogenetic tree constructed with 1,000 bootstrap replications via the neighbor joining method using MEGA 4.0 software (26).

### Determination of maximum tolerance concentration (MTC) for metals

The MTC of the isolates was determined by using the spot plate method with LB agar plates supplemented with CuSO_4_ concentrations ranging from 100 to 1000 mg/L, FeSO_4_ concentrations ranging from 100 to 1000 mg/L and MnSO_4_·4H_2_O concentrations ranging from 100 to 2000 mg/L (2, 3). The culture of 3 µL (corresponding to 0.5 OD at 600 nm) from each isolate grown overnight was spotted onto the plates and observed for visible growth after 3 d incubation at 30°С. The MTC was defined as the lowest concentration of metal without visible growth of the test strain.

### Investigation of effect of metals on bacterial growth

Isolate X1 that exhibited the highest MTC for 3 metals (Cu, Mn and Fe), isolate X26 that showed the highest MTC for Mn and the lowest MTC for Cu, and isolate X33 that displayed the highest MTC for Fe and the lowest MTC for Mn, were investigated for the effect of metals on their growth. The culture of 100 µL (corresponding to 0.5 OD at 600 nm) from the three strains grown overnight was transferred on to LB broth (100 mL) supplemented with middle concentrations of metals that the isolates could tolerate, i.e., isolate X1: 500 mg/L of CuSO_4_, 500 mg/L of FeSO_4_ and 1000 mg/L of MnSO_4_·4H_2_O; isolate X26: 300 mg/L of FeSO_4_ and 1000 mg/L of MnSO_4_·4H_2_O; isolate X33: 300 mg/L of CuSO_4_, 500 mg/L of FeSO_4_ and 1000 mg/L of MnSO_4_·4H_2_O. The media with bacteria but no metal were used as controls. The growth curves of the strains cultured on liquid medium with varied concentrations of metals at 30°C for 2 d with shaking at 150 rpm were produced by determining at OD_600_ using a SpectraMax iD3 microplate reader (Molecular Devices, CA, USA). All experiments were performed three times.

### Genome sequencing, assembly and prediction

Genomic DNAs were extracted from the isolates X1, X26 and X33 using GenElute bacterial genomic DNA extraction kit (Sigma-Aldrich, St. Louis, MO), and sequenced on a Nanopore platform (PromethION48) at Biomarker Technologies (Beijing, Chain). The filtered reads removed reads with mean_qscore_template of ˂ 7 and length of ˂ 2,000 bp from raw reads were assembled by Canu v1.5 software (27). Racon v3.4.3 software (28) was used to check the quality of the assembly, and then Circlator v1.5.5 software was used to cyclizing assembly genome (29). The prediction of coding genes and repetitive sequences was performed by Prodigal v2.6.3 (30) and RepeatMasker v4.0.5 (31), respectively. Transfer RNA (tRNA) and rRNA genes were predicted using tRNAscan-SE v2.0 (32) and Infernal v1.1.3 (33), respectively. PromPredict v1 and antiSMASH v5.0.0 were used for prediction of promoter (34) and secondary metabolic gene clusters (35), respectively.

### Genome annotation and comparation

For functional annotation, the protein sequences of the predicted coding sequences (CDSs) were blast against the databases, including Cluster of Orthologous Groups (COG), Gene Ontology (GO), Swiss-Prot, Kyoto Encyclopedia of Genes and Genomes (KEGG) and Non-Redundant Protein Database (NR). The subsystem analysis was performed by annotation using subsystems technology server (RAST) (36). To investigate the differences in metal adaptation strategies of isolates X1, X26 and X33, genes that are responsible for metal transport and metal binding proteins, which play a role in the metal adaptation in microorganisms, were searched from the genomes of the three strains and compared with each other. The genome sequences of isolates X1, X26 and X33 were deposited at Genbank under the accession number CP132290, JAUZEF000000000 and CP132291, respectively.

## ACKNOWLEDGMENTS

This research was supported by the funds from the “Hundred-Talent Program” of Guangdong Academy of Sciences (No. 2020GDASYL-20200102009 and 2020GDASYL-20200102010), the Pearl River Talent Plan Project (No. 2021CX020301), the GDAS’ Project of Science and Technology Development (No. 2022GDASZH-2022010110), and the Guangdong Basic and Applied Basic Research Foundation (No. 2022A1515110562), China.

